# A single motor neuron determines the rhythm of early motor behaviour in *Ciona*

**DOI:** 10.1101/2021.03.10.434256

**Authors:** Taichi Akahoshi, Kouhei Oonuma, Makoto Murakami, Takeo Horie, Takehiro G. Kusakabe, Kotaro Oka, Kohji Hotta

## Abstract

Vertebrate rhythmic motor behaviour is generated by the central pattern generator (CPG) located in the spinal cord. However, the development of the CPG has not been elucidated at the single-neuron level. We found that a single pair of motor neurons (A10.64/MN2) constitutes the CPG and regulates rhythmic early motor behaviour in the proto-chordate, *Ciona*. This pair of cells exhibited Ca^2+^ oscillation with an 80-sec interval at the mid-tailbud stage and 25 sec at the late tailbud stage. The Ca^2+^ oscillation occurred independently in a dissociated single cell. In the late tailbud stage, the Ca^2+^ oscillation began to coincide in phase with the ipsilateral tail muscle contraction, which corresponds to rhythmic early motor behaviour. Interestingly, the number and frequency of tail muscle contractions gradually coincided with spikes in the burst of membrane potential as the embryos developed toward late tailbud stage. Photoablation of A10.64/MN2 abolished the rhythmic early motor behaviour. These findings indicate that the early spontaneous rhythmic motor behaviour of *Ciona* is directly regulated by A10.64/MN2 as an essential component of the CPG. Our findings shed light on the development and evolution of chordate rhythmic behaviours.

## Introduction

Animal rhythmic motor behaviours are regulated by most, if not all, central pattern generators (CPGs). Although the neural circuits that constitute the CPG are not completely understood, they are defined by their ability to generate rhythmic motor patterns in the absence of sensory feedback or other rhythmic inputs ^1^. Since the historical study of Graham Brown, it has been believed that rhythmic motor behaviour in vertebrates is generated by a motor circuit located in the spinal cord ^2–4^.

During vertebrate early development, rhythmic spontaneous motor behaviour is observed before adult locomotion such as swimming, flying, and walking ^5–7^. Sporadic Ca^2+^ activities are first observed in the spinal cord, and then rhythmic spontaneous and patterned Ca^2+^ activities are gradually formed ^8–10^. However, the identification and development of neurons responsible for the chordate motor circuit is not fully understood, and it remains to be determined how the early motor circuit hands over control to the adult locomotor circuit.

Motor neurons have been traditionally considered the last relay from the central nervous system to the muscles. However, recent studies indicate that motor neurons may play important roles in rhythm generation for motor behaviour ^9,11,12^. On the other hand, interneurons called ipsilateral caudal neurons may also serve as pacemaker-like neurons that trigger rhythmic spontaneous motor behaviour ^13^. Thus, there is still no direct evidence regarding which neurons play the central role in pattern generation for rhythmic locomotor behaviour in chordates.

The chordate ascidian *Ciona* is the closest extant relative of the vertebrates ^14^. Early spontaneous motor behaviour in *Ciona*, termed “tail flick”, is first observed at the late tailbud stage ^15^. This behaviour is thought to facilitate hatching from the egg envelope ^15^.

Although a high-quality connectome of *Ciona* has been reported ^16^, the mechanisms and functions of the motor circuit controlling locomotion remain elusive.

In this study, to investigate the role of motor neurons in early spontaneous motor behaviour in *Ciona*, we performed Ca^2+^ and membrane potential imaging of neuronal activities, cell lineage tracking, single-cell ablation, and behavioural analysis. We found that a single motor neuron generates the rhythm for early spontaneous motor behaviour.

## Results

### Ca^2+^ oscillation in a pair of cells

As previously reported, oscillation of Ca^2+^ transients (Ca^2+^ oscillation) was observed symmetrically on the left and right sides of the motor ganglion (proposed to be homologous to the spinal cord of vertebrates) ^17^ (Fig. 1A), and no reproducible Ca^2+^ oscillation was observed in any other cells from the early-to mid-tailbud stage (St.20 to St.22; Fig. 1B white arrowhead, and Movie S1).

**Figure 1.**
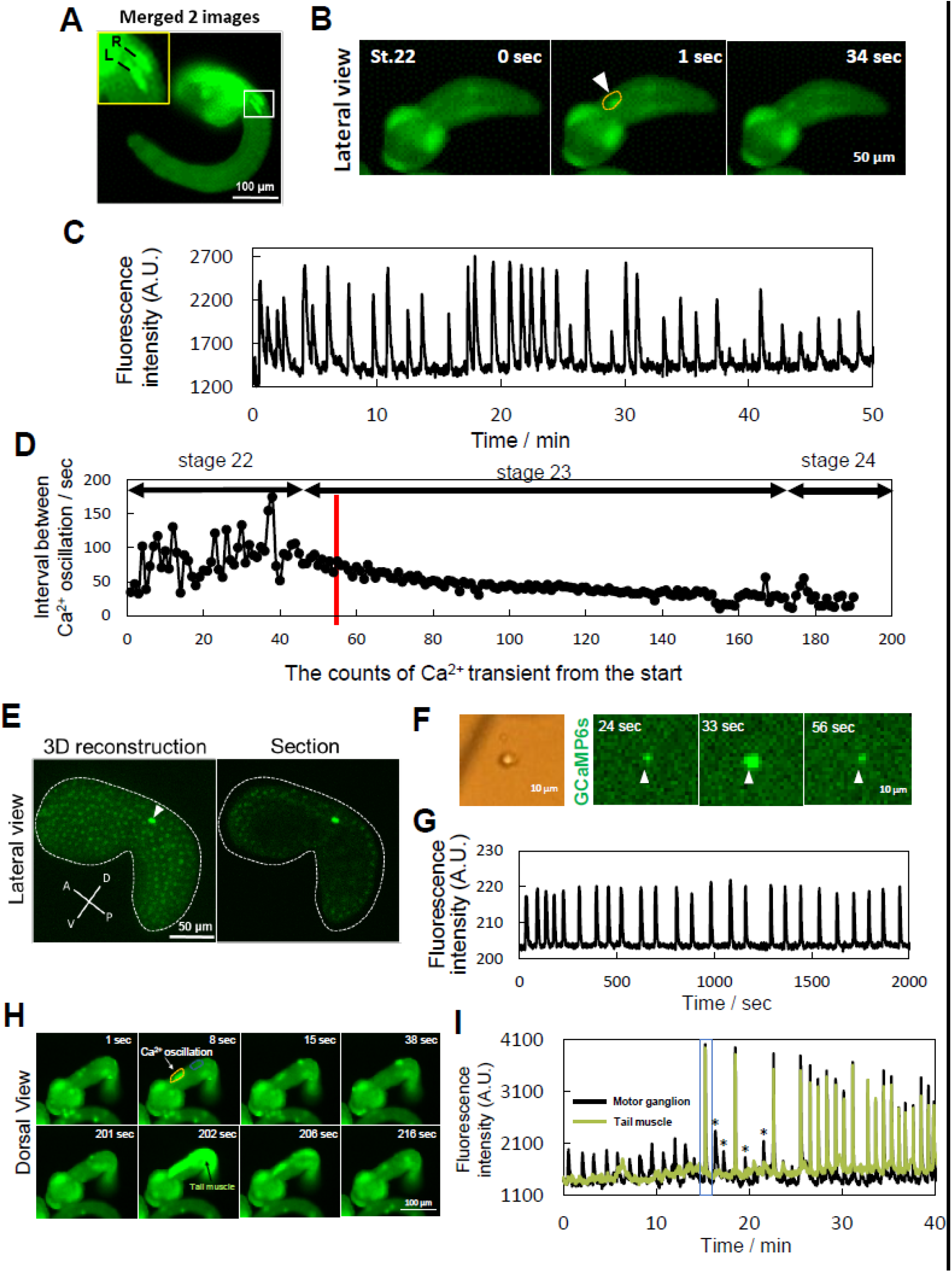
Ca^2+^ oscillation in a pair of cells synchronised with the activity of tail muscle cells at the tailbud stage. (A) Merged images of Ca^2+^ imaging at different time points. Ca^2+^ transients were observed on the left and right sides of the motor ganglion, respectively (yellow box; enlarged view of the white box, inset). R, right; L, left. (B) Representative time course image of the Ca^2+^ transient in the motor ganglion at St.22. Ca^2+^ transient is indicated by a white arrowhead (second panel). (C) Fluorescence intensity of Ca^2+^ in the ROI (region of interest, Fig.1B, yellow dotted line) over 50 min. (D) Transition of the interval between Ca^2+^ oscillations over time. As the count of Ca^2+^ transients increases, the interval between Ca^2+^ oscillations gradually decreases. Red line indicates the time when Ca^2+^ oscillations first coincided with the Ca^2+^ elevation of the ipsilateral muscle. Approximate developmental stage is indicated. (E) 3D reconstruction (left panel) and section (right panel) images of the Ca^2+^ oscillation detected by H2B-GCaMP6s. Ca^2+^ oscillation is indicated by the white arrowhead. Ca^2+^ oscillation occurs only in a single nucleus. A, anterior; D, dorsal; P, posterior; V, ventral. (F) Image of a dissociated single cell and time-course images of Ca^2+^ oscillation in the same cell. N = 6. (G) Change in fluorescence intensity of Ca^2+^ in (F) over 2000 sec. (H) Representative time-course image of Ca^2+^ oscillation at St.23. After a few minutes, the Ca^2+^ oscillation (white arrow, 8 sec) coincides with the Ca^2+^ elevation in the ipsilateral muscle cells (black arrow, 202 sec). (I) Fluorescence intensity over time in the motor ganglion and tail muscle at St.23. ROIs: Fig.1H, 8 sec, yellow dotted line for motor ganglion, blue dotted line for tail muscle cell. The time when the Ca^2+^ oscillation and Ca^2+^ elevation of ipsilateral muscle cells first coincided is indicated by a blue rectangle. The asterisks indicate Ca^2+^ oscillation without synchronisation of Ca^2+^ elevation in ipsilateral muscle cells after the first synchronisation.

In mid-tailbud II (St.22), the duration and interval of the Ca^2+^ oscillation on one side were 23 ± 4 (mean ± S.D.) sec and 80 ± 4 (mean ± S.D.) sec, respectively (Fig. 1C, 1D). The interval of Ca^2+^ oscillations varied at St.22, but gradually decreased to 25 sec toward late tailbud II (St.24) (Fig. 1D). The duration of Ca^2+^ oscillation also gradually decreased [12 ± 2 (mean ± S.D.) sec at St.23 and 10 ± 2 (mean ± S.D.) sec at St.24].

To determine how many cells are involved in the Ca^2+^ oscillation, we expressed the nuclear-localised Ca^2+^ sensor H2B-GCaMP6s in the whole embryo. Time-lapse imaging of the lateral side by confocal microscopy revealed that Ca^2+^ oscillation occurred only in a single nucleus (Fig. 1E and Movie S2), and no other cells exhibited reproducible Ca^2+^ transients at mid-tailbud II (St.22), indicating that the Ca^2+^ oscillation was derived from only a single pair of cells at mid-tailbud II (St.22).

### A cell-autonomously oscillating neuron

To determine whether a single cell has the potential to exhibit the Ca^2+^ oscillation independently, we dissociated tailbud embryos expressing GCaMP6s into single cells and observed Ca^2+^ levels (Fig. 1F and Movie S3). Surprisingly, time-lapse imaging revealed a cell-autonomous Ca^2+^ oscillation at an interval of 40–80 sec (Fig. 1G). Thus, the Ca^2+^ oscillation during the tailbud stage was caused by a single cell, acting independently without any contribution from other cells.

Continuous whole-embryo Ca^2+^ imaging revealed that the Ca^2+^ oscillation began at the same time as Ca^2+^ elevation in the ipsilateral tail muscle from late tailbud I (St.23). Earlier Ca^2+^ oscillations in the motor ganglion sometimes did not elevate Ca^2+^ in the tail muscle (Fig. 1I asterisks), but fully synchronised 10 min after the first synchronisation (Fig. 1H at 202 sec and Fig. 1I, blue box). Tail muscle contraction was also observed at the same time as Ca^2+^ elevation in the tail muscle (at 7 min in Movie S4). These results suggest that the pair of neurons exhibiting Ca^2+^ oscillation are motor neurons, and that these cells directly regulate the contractions of tail muscle after late tailbud I (St.23).

We investigated the cell lineage of a pair of cells exhibiting Ca^2+^ oscillation. We expected that a largest motor neurons A10.64 corresponded to this pair of cells (Fig. 2A). Simultaneous imaging of Neurog::Kaede-NLS and pSP-CiVACHT::GCaMP6s can track the cell lineage of A9.31 and A9.32 (mother cell of A10.64; Fig. 2B and 2C). A pair of Kaede nucleus-localised signals of A10.64 appeared exclusively in the motor ganglion at late tailbud I (St.23) (Fig. 2D, 137 min). Finally, we confirmed that the Ca^2+^ oscillation in the cytoplasm overlapped with the Kaede nucleus–localised signal of A10.64 (Fig. 2E yellow arrowhead, Movie S5). Based on these findings, we identified the cells exhibiting Ca^2+^ oscillation as A10.64, which are MN2 cholinergic motor neurons ^16,18^.

**Figure 2.**
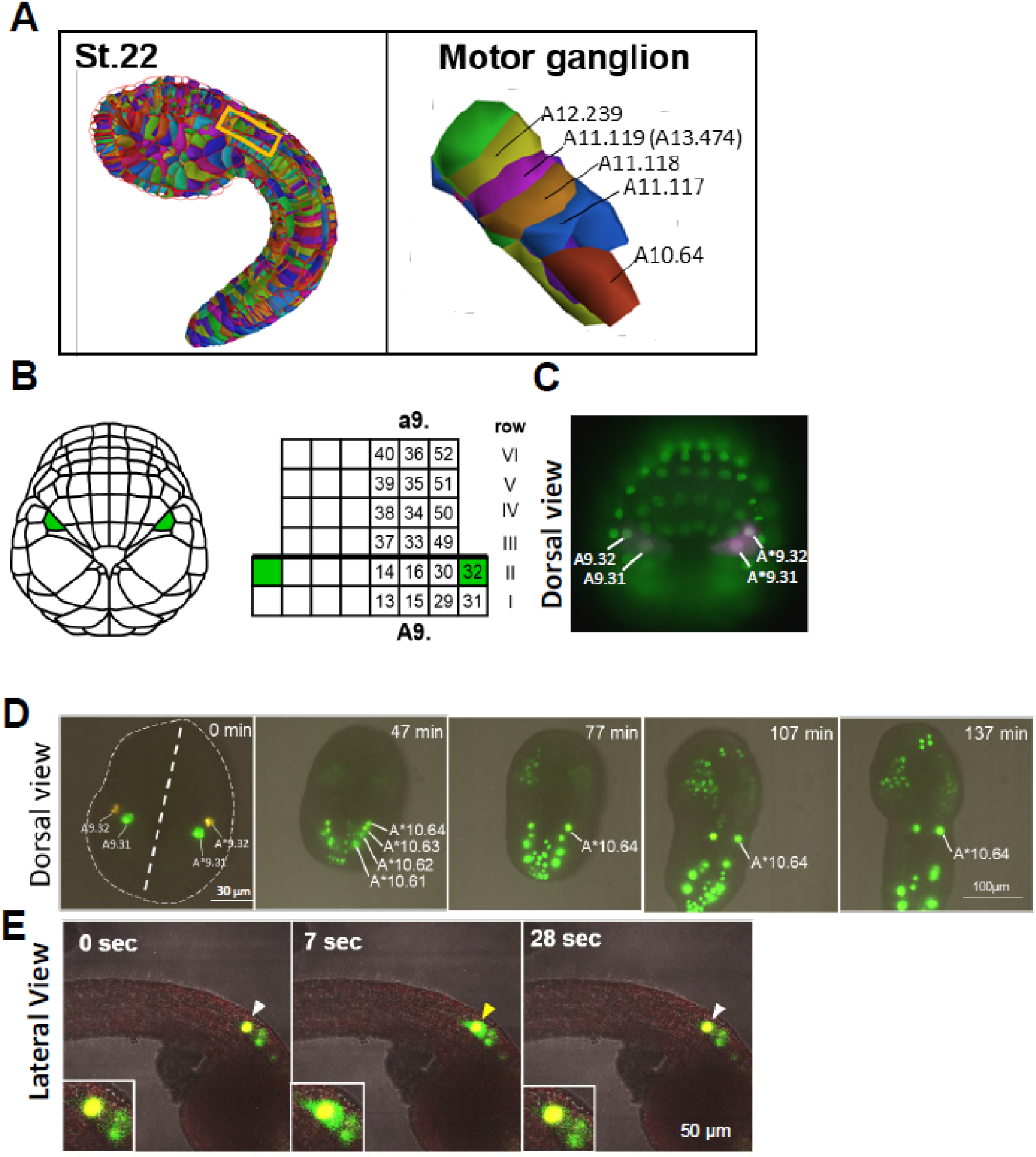
The motor neuron MN2/A10.64 causes cell-autonomous Ca^2+^ oscillation. (A) 3D reconstruction of a *Ciona* embryo at mid-tailbud II (St.22), modified from ^32^. The region of the motor ganglion (yellow rectangle) is magnified in the right panel. The cell lineage of five pairs of cholinergic neurons is indicated. A11.119 is the mother cell of the motor neuron A13.474. (B) Schematic illustrations of the late gastrula (St.13) (left) and the cell lineage of the neural plate cells (right). A9.32 cells are indicated in green. (C) Late gastrula embryo expressing Kaede under the control of the Neurogenin promoter. Kaede was immunostained (magenta). Nuclei were stained with DAPI (green). In reference to Fig. 2A, A9.32 and A9.31 cells are identified. A9.32 and A9.31 cells overlapped with Kaede signals. (D) Cell lineage tracking of A9.32 cells from late gastrula (St.13) to late tailbud I (St.23). The embryo was electroporated with Neurog::Kaede-NLS and pSP-CiVACHT::GCaMP6s. The sister cell of A9.32, A10.64, migrates anteriorly and is localised to the motor ganglion at late tailbud I (St.23). (E) Representative time-course images of Ca^2+^ oscillation at St.23. A10.64 is labelled with Kaede-NLS (white arrowhead). Ca^2+^ oscillation overlapped with the nuclear signal from Kaede-NLS (yellow arrowhead). Enlarged views of A10.64 are embedded in each panel (inset). N = 6.

### Muscle contraction and spikes of A10.64

To understand the relationship between Ca^2+^ oscillation and membrane potential, we performed simultaneous recordings of Ca^2+^ and membrane potential using the red fluorescent Ca^2+^ indicator NES-jRGECO1a ^19^ and the voltage sensor ASAP2f ^20^. In the Ca^2+^ transient (Fig. 3A and Fig. S1, red graph), we could observe burst firing of membrane potential in A10.64 (Fig. 3A and Fig. S1, green graph). Each burst consisted of multiple spikes (Fig. 3A and Fig. S1, asterisks). Because every burst was accompanied by a single Ca^2+^ transient, the timing and interval of the bursts were consistent with those of the Ca^2+^ oscillations (Fig. 1E).

**Figure 3.**
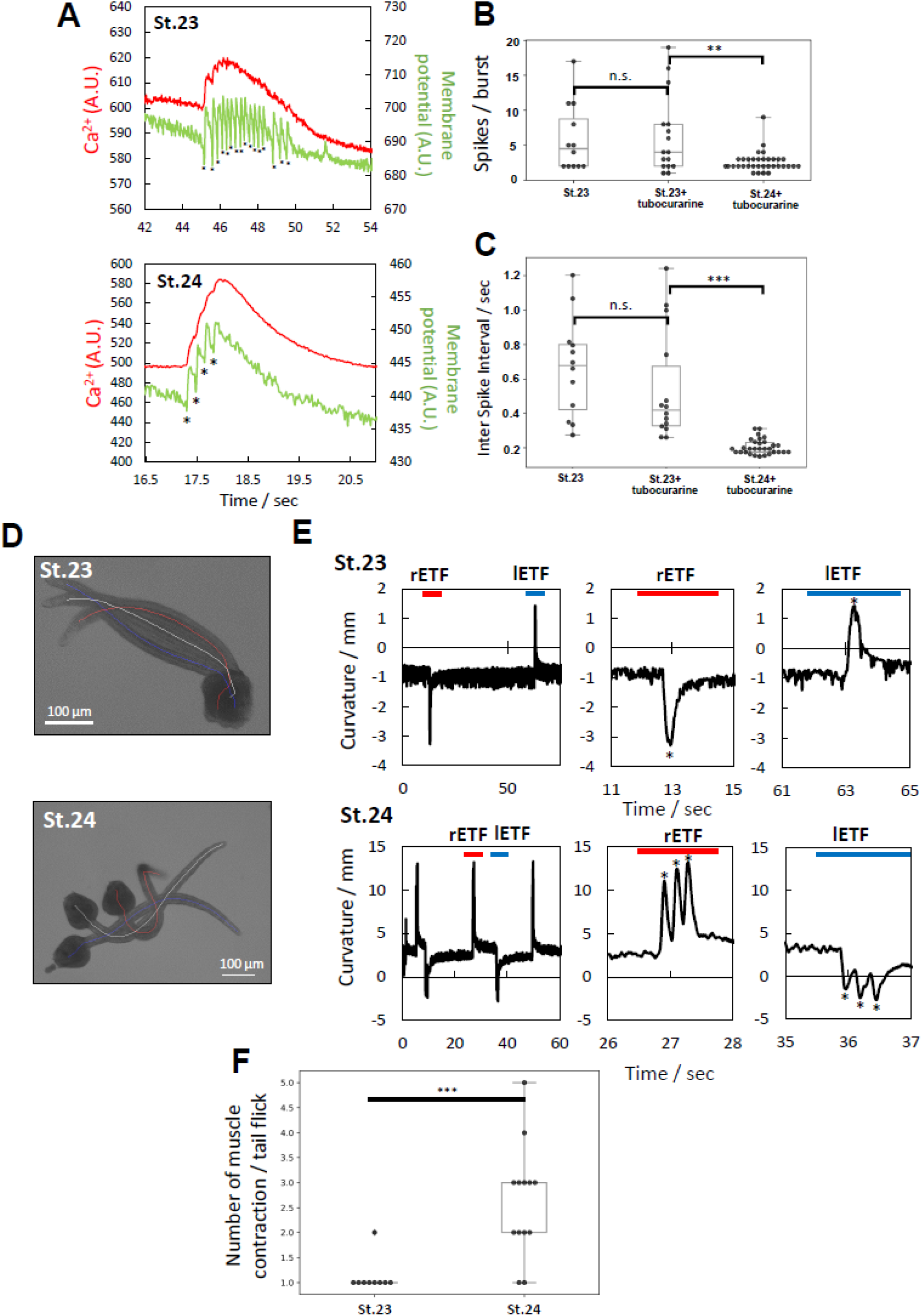
Muscle contraction couples with the neuronal activity of A10.64. (A) Representative fluorescence intensity of Ca^2+^ (red) and membrane potential (green) in A10.64 at St.23 (top panel) and St.24 (lower panel). The fluorescence intensity of ASAP2f decreases upon depolarisation ^20^. Asterisks indicate the timing of depolarisation. (B) Box plot of numbers of spikes in bursts in A10.64. Data are presented as means ± S.D. (n = 7, 12, and 11 for tubocurarine 0 mM (St.23), 5 mM (St.23), and 5 mM (St.24), respectively; ** P < 0.05; Wilcoxon rank–sum test). (C) Box plot of inter-spike intervals in A10.64. Data are presented as means ± S.D. (n = 7, 11, and 11 for tubocurarine 0 mM (St.23), 5 mM (St.23), and 5 mM (St.24), respectively; ** P < 0.05; Wilcoxon rank–sum test). (D) Three merged images of late tailbud I (St.23, upper panel) and late tailbud II (St.24 lower panel), captured by a high-speed camera. The midlines of the embryos are indicated in white (resting state), red (timing of right muscle contraction), and blue (timing of left muscle contraction). (E) Time course of curvature, calculated from the midline at late tailbud I (St.23, upper panel) and late tailbud II (St.24 lower panel). Red and blue bars indicate representative right and left muscle contractions, respectively. Magnified time course of curvature at red and blue bars is indicated in the right graphs, respectively. Asterisks indicate the timing of tail muscle contraction. (F) Box plot of the number of muscle contractions in one ETF at St.23 and St.24. Data are presented as means ± S.D. [n = 9 (St.23) and 13 (St.24) from 3 and 2 embryos respectively; *** P < 0.01; Wilcoxon rank–sum test].

The average number of spikes in a burst was 6.3 ± 5.4 at late tailbud I (St.23), decreasing significantly to 2.6 ± 1.4 at late tailbud II (St.24) (p < 0.05) (Fig. 3B). The inter-spike interval in a burst was 0.5 ± 0.3 sec at late tailbud I (St.23), decreasing significantly to 0.2 ± 0.04 sec at late tailbud II (St.24) (p < 0.05) (Fig. 3C).

Tail muscle contraction was first observed in St.23, when Ca^2+^ elevation occurred in the tail muscle (Movie S4). This suggested that in St.23, Ca^2+^ oscillations in the A10.64 elevate Ca^2+^ levels in the tail muscle and fully synchronise the muscle contractions (Fig. 1I and 1J). In that case, what about the relationship between the neuronal activity of A10.64 and muscle contractions at stages after St.23? Because of its unique function, A10.64/MN2 are expected to regulate perpetual rhythmic behaviour in later stages as well.

Considering that the interval of the Ca^2+^ oscillation in A10.64 gradually decreased from 80 sec (St.22) to 25 sec (St.24) (Fig. 1D), and tail muscle contraction was coupled with A10.64 after late tailbud I (St.23), we expected that the serial muscle contractions would occur every 80 sec (St.22) and 25 sec (St.24), respectively. To test this, we used a high-speed camera to record the muscle contraction both on the left and right sides at St.23 and St.24 (Fig. 3D and 3E). We referred to each set of unilateral muscle contraction as an early tail-flick (ETF). The average interval between ETFs was 79 ± 14 (mean ± S.D.) sec at St.23, decreasing to 21 ± 6 (mean ± S.D.) sec at St.24, comparable to the change in the Ca^2+^ oscillation in A10.64 between St.23 and St.24 (80 sec to 25 sec; Fig. 1E).

Interestingly, tail muscle contraction occurred only once in each ETF at St.23, whereas two or three consecutive muscle contractions occurred in each ETF at St.24 (Fig. 3E). The number and frequency of tail muscle contractions in each ETF at St.24 (Fig. 3E, F) were comparable to those of spikes in the burst at St.24 (Fig. 3B and C). These observations indicated that muscle contractions in each ETF gradually coupled to the individual spike in the burst of A10.64 as the embryo developed toward St.24.

### Photoablation of A10.64

We hypothesised that the neuronal activity of the A10.64 would be sufficient to regulate the ipsilateral ETF at least until St.24. To test this hypothesis, we performed single-cell photoablation of A10.64 using an LOV-based optogenetic tool, miniSOG2, which produces singlet oxygen under laser irradiation ^21^. In contrast to the control embryo (electroporated only with Neurog::mCherry-CAAX), the embryo expressing miniSOG2 did not exhibit ETFs at St.24 (Fig. 4C and 4D). In the absence of 440-nm laser irradiation, early motor behaviour was comparable to that in control embryos (Fig. S2).

**Figure 4.**
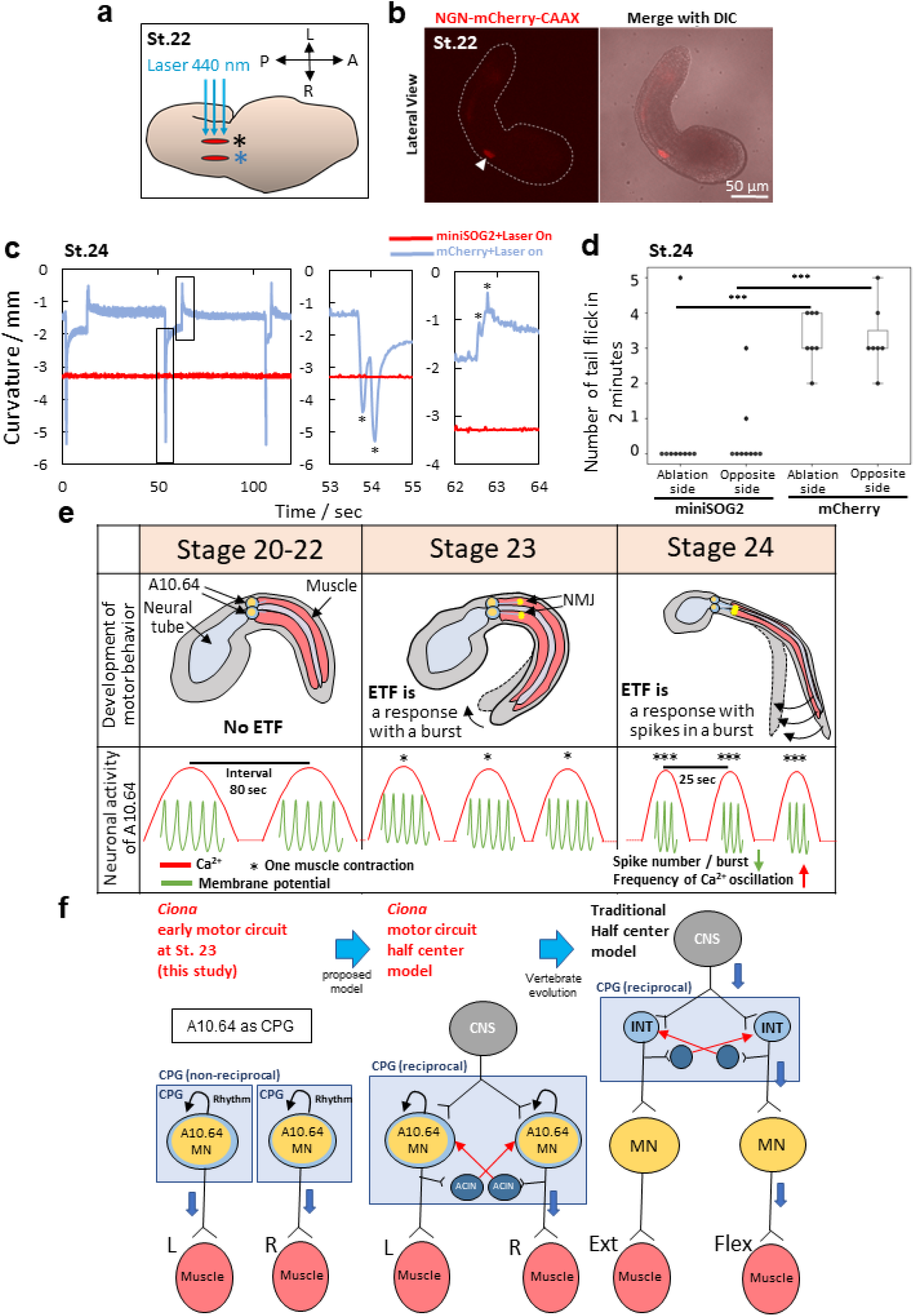
Single-cell photoablation of A10.64 abrogates ETFs until St.24. (A) Schematic illustrations of photoablation of A10.64. A10.64 cells are shown in red. A 440-nm laser was applied from the lateral side. The upper and lower sides of A10.64 are marked by black and blue asterisks, respectively. (B) Fluorescence image (left) and merged image with DIC (right) at mid-tailbud II (St.22) in an embryo electroporated with Neurog::mCherry-CAAX. The signal of mCherry-CAAX in A10.64 is indicated by a white arrowhead. (C) Time course of curvature, calculated from the midline, for 120 sec. Blue line indicates the curvature of embryos electroporated with Neurog::mCherry-CAAX (for control). Red line indicates the curvature of embryos electroporated with Neurog::miniSOG2-CAAX and Neurog::mCherry-CAAX. Magnified time course of curvature (black rectangles) is shown in the right graphs. (D) The number of ETFs in 2 min at St.24 under each condition. The upper side of A10.64 (Fig. 5A, black asterisk) is indicated as the “Ablation side”. The lower side of A10.64 (Fig. 5A, blue asterisk) is indicated as the “Opposite side”. Data are presented as means ± S.D. [N = 9 (electroporated with Neurog::miniSOG2-CAAX and Neurog::mCherry-CAAX) and N = 7 (electroporated with mCherry-CAAX, for control); *** P < 0.01; Wilcoxon rank–sum test]. (E) Summary of the Ca^2+^ oscillation and membrane potential in A10.64, and its relationship to the early spontaneous motor behaviour from St.22 to St.24. Neural tubes (blue), muscle (red), A10.64 (green), and neuromuscular junction (NMJ, yellow) are indicated by black arrows. The asterisks and curved arrow indicate the number of muscle contractions in one Ca^2+^ oscillation or ETF. (F) (left panel) Schematic of *Ciona* early motor circuit at St. 23 identified in this study, showing that a single motor neuron generates the rhythm of motor behaviour and directly controls muscle contraction. (middle) Schematic of *Ciona* early motor circuit. Independent CPGs in both left and right sides in St. 23 become one reciprocal CPG by joining ACIN commissure neurons ^33^. (right) Traditional half center model, in which the central pattern generator (CPG) consists of interneurons that drive activity in motor neurons; in turn, motor neurons relay these patterns to muscles, which produce behaviour ^11^.

Together, these results suggested that oscillatory neuronal activity of a single pair of cells, A10.64, directly regulates ETFs until St.24, which means that the spontaneous oscillatory firing of A10.64 motor neuron directly commands rhythmical tail muscle contraction. Although single-cell photoablation of A10.64 abolished early spontaneous motor behaviour until St.24, tail muscle contraction recovered at a later stage (St. 26, Movie S6). These observations suggest that another type of motor neuron starts to be active and forms a new neuromuscular junction after St.24 ^22^.

## Discussion

### Development of ascidian motor circuit

In this study, we report direct evidence of a single pair of motor neurons, A10.64, that play a central role in pattern generation of early rhythmic locomotor behaviour in *Ciona*. Specifically, we found that A10.64 exhibits Ca^2+^ oscillation with an 80-sec interval in St.22 (Fig. 1C, Fig.1D, Fig.2E), and that A10.64 oscillates independently (Fig. 1F, G). Furthermore, we revealed the relationship between the Ca^2+^ oscillation, the burst firing, and spikes of membrane potential in the early spontaneous motor behaviour. Based on our results, early development of *Ciona* motor neural circuit proceeds as follows (Fig. 4E).

1. From early to mid-tailbud II (St.20 to St.22), coupling of Ca^2+^ oscillation with burst firing of membrane potential first occurs only in A10.64 (Fig. 1E). The interval and the duration of the Ca^2+^ oscillation are approximately 80 sec and 23 sec, respectively (Fig. 1D). No muscle contraction is observed in this stage.
2. In late tailbud I (St.23), A10.64 first coincides with the Ca^2+^ elevation of ipsilateral tail muscle (ETF first observed), suggesting the formation of neuromuscular junctions (Fig. 4E, yellow stars). One tail muscle contraction coupled with one burst firing is associated with multiple spikes.
3. In late tailbud II (St.24), both the interval and duration of each Ca^2+^ transient are shorter, approaching 25 sec and less than 10 sec, respectively (Fig. 1D, Fig. 3A). In the ETF of St. 24, each tail muscle contraction coincides with each spike in a burst.

In this manner, the early spontaneous motor behaviour of *Ciona* begins with the activation of the oscillatory motor neuron A10.64; toward later stages, muscle contraction of ETF is gradually fine-tuned to each spike, rather than each burst of membrane potential in A10.64 (Fig. 4E).

### Roles of A10.64 motor neuron as CPG

Our results suggest that the motor neuron A10.64 has the function similar to that of CPG ^1^ or a rhythm-generating pacemaker-like neuron ^23^ (Fig.4F). CPGs are experimentally robust because they can often be activated and studied in isolated brains, producing ‘fictive behaviours’ in which circuit output closely resembles naturally observed behaviour patterns ^24^.

Supporting this, A10.64 is the sole oscillator at St.22 (Fig.1E), and isolated A10.64 oscillates independently in a manner that closely resembles naturally observed rhythms of ETFs (Fig. 1G) and synchronises with muscle activity (Fig.1 H, I). Furthermore, inhibition of the oscillation halts rhythmic ETFs (Fig. 4C, D). Hara et al. (2021 preprint)^25^ recently reported that decerebrated *Ciona* larvae that contain motor ganglion region exhibit tail beating bursts with an interval of approximately 20 sec, similar to the interval of the Ca^2+^ oscillation in A10.64 at St. 24 (Fig. 1E).

### Role of A10.64 motor neuron in larva

Ascidian matured larvae spontaneously exhibit swimming-like alternating left and right movements starting from St.26 ^26^, but ETFs in St.23 to St.24 are not coupled between the left and the right side. How does ETF behaviour in early stages transition to alternating swimming behaviour in the mature larva?

In principle, the motor circuit involved in locomotion consists of different components, including a command neuron that receives input from sensory afferent and is necessary and sufficient for initiating a variety of behavioural responses ^27,28^. The CPG typically requires the input from command functions in order to generate reciprocal rhythmic neuronal activity ^29^.

The “half center” hypothesis proposes that rhythmic motor activity is generated by reciprocal inhibition between two pools of interneurons located on each side of the spinal cord—an extensor half center activating extensor motoneurons and a flexor half center exciting flexor motoneurons ^30^.

Ascidian larva are also thought to achieve left–right alternating muscle contraction by adding reciprocal connections from anterior caudal inhibitory neurons (ACINs) to the motor circuit (Fig. 4F ^31^). ACINs are interneurons that extend axons anteriorly to connect to contralateral motor neurons ^16^. Given that A10.64 will continue to oscillate permanently (Fig.1G), it is possible that spontaneous motor command by A10.64 continues through later larval stages, generating a reciprocal rhythmic neuronal activity as a component of the reciprocal inhibitory system after St.24.

A widespread mechanism that contributes to rhythm generation in CPGs is post-inhibitory rebound, in which a neuron will reliably spike following inhibitory input from another neuron (or group of neurons) in the circuit ^1^. The *Ciona* interval of left–right muscle contraction changed from a few seconds in the tailbud stages (Fig.3E) to 15–25 ms in mature swimming behaviour of hatched larvae ^26^. The shortened interval between left and right muscle contractions might be the result of the post-inhibitory rebound between the left and right side of A10.64s due to joining the reciprocal inhibitory system of the ACINs (Fig. 4F). Thus, the A10.64 motor neuron, which has special characteristics, is considered to be an essential component of mature larval locomotion circuits.

## Supporting information

**Suppl. Figure 1.**
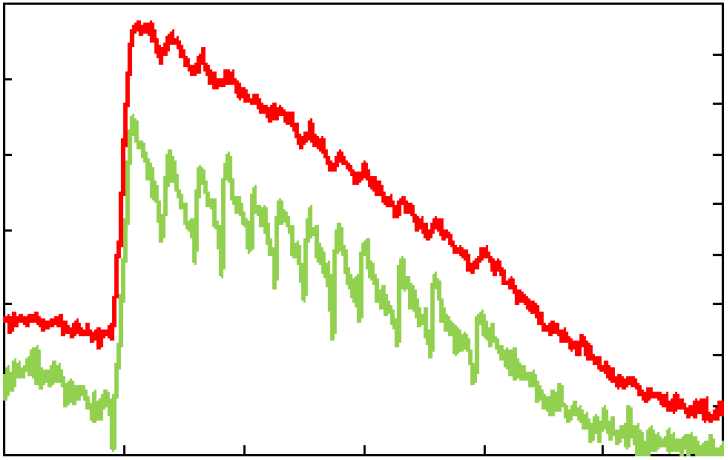
Representative fluorescence intensity of Ca^2+^ and membrane potential in A10.64 at St.22. Fluorescence intensity of Ca^2+^ and membrane potential are indicated in red and green, respectively. Fluorescence intensity of ASAP2f decreases upon depolarisation ^34^. Asterisks indicate the timing of depolarisation.

**Suppl. Figure 2.**
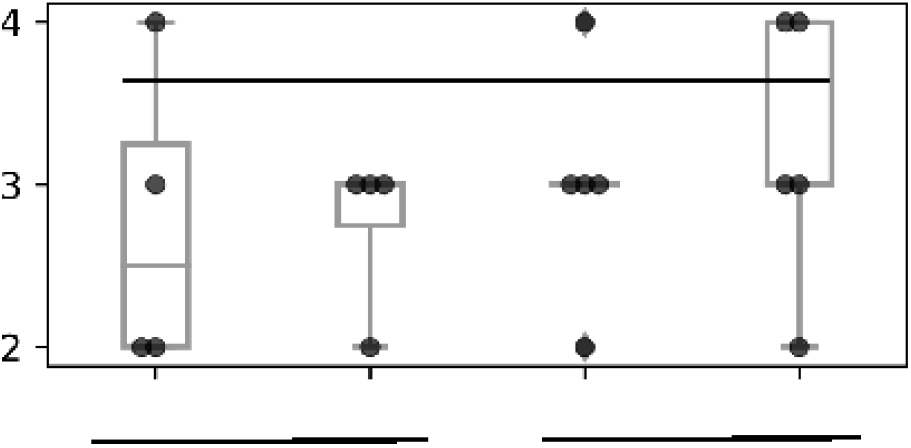
Number of ETFs observed in 2 min at St.24 under each condition. Embryos were electroporated with Neurog::miniSOG2-CAAX and Neurog::mCherry-CAAX (indicated as miniSOG2), or Neurog::mCherry-CAAX (indicated as mCherry). No laser irradiation was applied to A10.64 under any condition. Data are presented as means ± S.D. [N = 4 (electroporated with Neurog::miniSOG2-CAAX and Neurog::mCherry-CAAX) and N = 5 (electroporated with mCherry-CAAX, for control); n.s., not significant; Wilcoxon rank–sum test].

**Suppl. Movie 1. Whole-body Ca**^**2+**^ **imaging using GCaMP6s at mid-tailbud II (St.22)**.

Fluorescence imaging of mid-tailbud II (St.22) expressing GCaMP6s. Ca^2+^ oscillation was observed in the motor ganglion. The other Ca^2+^ transients observed in this video were derived from epidermis ^35^ and did not couple with the Ca^2+^ oscillation in the motor ganglion. Time-lapse imaging was recorded from the lateral side.

**Suppl. Movie 2. Whole-body Ca**^**2+**^ **imaging using H2B-GCaMP6s at mid-tailbud II (St.22)**.

Fluorescence imaging of mid-tailbud II (St.22) expressing H2B-GCaMP6s, recorded by confocal laser-scanning microscopy. Ca^2+^ oscillation was observed only in a single nucleus in the motor ganglion. The other Ca^2+^ transients observed in this video were derived from the epidermis ^35^ and did not couple with the Ca^2+^ oscillation in the motor ganglion. The time-lapse images were recorded from the lateral side.

**Suppl. Movie 3. Ca**^**2+**^ **oscillation in a dissociated single cell**

Imaging of the dissociated single cells expressing GCaMP6s was performed by fluorescence microscopy. Time-lapse images were recorded at 1-sec intervals.

**Suppl. Movie 4. Long time-lapse Ca**^**2+**^ **imaging using GCaMP6s, from mid-tailbud II (St.22) to late tailbud II (St.24)**.

Fluorescence imaging of embryos expressing GCaMP6s. After about 7 min, the Ca^2+^ elevation in the tail muscle gradually came to coincide with the ipsilateral tail muscle contraction. The Ca^2+^ transients derived from epidermis are also observed in this video ^35^.

**Suppl. Movie 5. Simultaneous imaging of Neurog::Kaede-NLS and pSP-CiVACHT::GCAMP6s at late tailbud I (St.23)**.

Simultaneous imaging of Neurog::Kaede-NLS and pSP-CiVACHT::GCAMP6s at late tailbud I (St.23), recorded by confocal laser-scanning microscopy. A10.64 labelled with Kaede-NLS exhibited Ca^2+^ oscillation. Time-lapse images were recorded from the lateral side.

**Suppl. Movie 6. Bright-field recording of a hatched larva (St.26) in which A10.64 was laser-ablated at St.22**.

In the hatched larva (St.26) shown in this video, A10.64 was ablated at St.22. The embryo did not exhibit early spontaneous motor behaviour at St.24 (Fig. 4C, red lines). Muscle contraction occurred frequently at St.26, although normal swimming behaviour was not observed.

## Methods

### Experimental Animals

*Ciona robusta* (*Ciona intestinalis* type A) adults were obtained from Maizuru Fisheries Research Station (Kyoto University), Onagawa Field Center (Tohoku University), and Misaki Marine Biological Station (The University of Tokyo) through the National Bio-Resource Project (NBRP), Japan. Eggs and sperm were collected by dissection of oviducts and sperm ducts, respectively. After artificial insemination, fertilised eggs were incubated at 15–20°C until observation.

### Preparation of reporter constructs and gene transduction

pGP-CMV::GCaMP6s ^36^, pGP-CMV::NES-jRGECO1a ^19^, ASAP2f ^20^, and pcDNA3.1::miniSOG2-T2A-H2B-EGFP ^21^ were purchased from Addgene (USA). pSP-CiVACHT::Kaede was purchased from *<u>C</u>iona intestinalis* <u>T</u>ransgenic line <u>RES</u>ources (CITRES). Neurog::Kaede-NLS was made as follows: a 3.6-kb upstream region of the *Ciona neurogenin* gene (*Neurog*) was amplified by PCR using a pair of primers (forward primer, AGGGATCCGGAAGAGGTGTTAGA; reverse primer, GGGGATCCATTTTGTAGCAAGAGC) and cloned into vector pSP-Kaede-NLS vector ^37^. pSP-Neurog::Kaede, which was used for immunostaining of Kaede at late gastrula, was made as follows: a 3.6-kb upstream region of the *Ciona neurogenin* gene (*Neurog*) was subcloned into the *Bam*HI restriction site of the pSP-Kaede vector.

Simultaneous imaging of Neurog::Kaede-NLS and pSP-CiVACHT::GCaMP6s can track the cell lineage of A9.31 and A9.32 (mother cell of A10.64; Fig. 2B and 2C) as well as observe the Ca^2+^ activity in the cytoplasm of pan-cholinergic motor neurons ^34^. Time-lapse recording starting in late gastrula revealed that A9.32 divided at mid-neurula, and that A10.64 migrated anteriorly along the neural tube (Fig. 2D), as reported previously ^38^.

For micro-injection, mRNA encoding GCaMP6s was synthesised from pSPE3::GCaMp6s as previously described ^35^. By injecting mRNA encoding GCaMP6s into unfertilised eggs, we could observe Ca^2+^ transients in the whole embryo from early gastrula (St.11) to late tailbud III (St.25) 24. For synthesis of mRNA encoding H2B-GCaMP6s, the ORF of H2B was PCR-amplified from Foxb::H2B-CFP (forward primer, TCTGAATTCAGGCCTATGGTTGCATCCAAA; reverse primer, GGCGACTGGTGGATCTTTTGAGCTGGTGTA), and GCaMP6s was PCR-amplified from pSPE3::GCaMp6s (forward primer, GATCCACCAGTCGCCACCATGGGTTCTCAT; reverse primer, AGGCCTGAATTCAGATCTGCCAAAGTTGAG). The cloning reaction for pSPE3::H2B-GCaMp6s was performed using linearised PCR products and the In-Fusion HD Cloning Kit (Takara Bio). H2B-GCaMP6s mRNA was produced and precipitated using the mMESSAGE mMACHINE T3 kit (Life Technologies, Carlsbad, CA, USA). GCaMP6s or H2B-GCAMP6s mRNA was injected into dechorionated eggs at 0.5 μg/μl.

For electroporation, pSP-CiVACHT::GCaMP6s, pSP-Neurog::ASAP2f, pSP-Neurog::NES-jRGECO1a, pSP-Neurog::miniSOG2-CAAX, and pSP-Neurog::mCherry-CAAX were synthesised as described for pSPE3::H2B-GCaMp6s. Forty microliters of each plasmid-construct at 1000 ng/μl was combined with 360 μl of 0.77 M mannitol in 10% seawater. At 30 min post-fertilisation, eggs in 400 μl of this solution were placed in a cuvette for electroporation. After electroporation, eggs were washed with seawater and incubated until observation. For membrane potential imaging, embryos at St.23 and St.24 were paralysed with 5 mM D-tubocurarine 30 min before observation, as tail muscle contraction prevents measurement of membrane potential ^39^. D-tubocurarine affected neither the number of spikes in the burst nor the inter-spike interval (ISI) in embryos at St.23 (Fig. 3B, 3C).

### Immunostaining of late gastrula embryo with Kaede antibody and DAPI

Immunofluorescence staining was carried out as described previously ^33^. To visualise the localisation of fluorescent reporter proteins, polyclonal rabbit anti-Kaede (PM012; Medical & Biological Laboratories, Nagoya, Japan; for Kaede) was used as the primary antibody (diluted 1:1000). The secondary antibody was Alexa Fluor 594–conjugated anti-rabbit IgG (A11012; Thermo Fisher Scientific). Samples were mounted under a coverslip in 50% glycerol-PBST with mounting Medium with DAPI (Vector Laboratories).

### Microscopy

The embryos were observed by fluorescence microscopy with a 3CCD (C7800-20, Hamamatsu Photonics) camera or by confocal laser-scanning microscopy (CLSM). A Nikon inverted microscope (Nikon eclipse, IX71) with a 20× or 40× objective lens (LUCPlanFLN) was used for fluorescence imaging with a U-MWBV2 mirror unit (Olympus). A SOLA LED light (Lumencor) was used as the light source, and fluorescence images were acquired with a 3CCD camera and the AQUACOSMOS software (Hamamatsu Photonics). For Ca^2+^ imaging, the time interval was set to 1–5 sec per frame. For membrane potential imaging, the time interval was set to 20 msec per frame. For CLSM imaging, an Olympus fv1000 microscope was used. Excitation was performed at 488 nm to visualise the signals of GCaMP6s and Kaede, and at 559 nm to visualise the signal of mCherry. An Olympus 20× or 40× oil immersion lens was used.

### Fluorescence image analysis

Fluorescence intensities in image data were analysed using the AQUACOSMOS software. Numerical data were exported into Microsoft Office Excel 2016 (Microsoft, Redmond, WA, USA) for graph plotting. Statistical analyses were performed in R (CRAN).

### Cell lineage tracking

The Z-stack images of the dorsal side of late gastrula embryos electroporated with Neurog::Kaede-NLS and pSP-CiVACHT::GCaMP6s were acquired every 10 min until late tailbud I (St.23) using an Olympus fv1000 microscope. The photoconversion of Kaede was performed by bleaching with a SIM scanner by manual operation. The Kaede signal of a pair of A9.32 cells at late gastrula (St.13) was photoconverted by irradiation with a 405-nm laser for a few seconds.

### Single-cell photoablation of A10.64

Single-cell photoablation of A10.64 was performed using miniSOG2, which produces singlet oxygen by laser irradiation. pSP-Neurog::miniSOG2-CAAX was used to express miniSOG2 at A10.64, and pSP-Neurog::mCherry-CAAX was used to label the A10.64 for laser irradiation.

In mid-tailbud II (St.22), mCherry-CAAX was expressed exclusively in A10.64 (Fig. 4B). Embryos (St.22) electroporated with pSP-Neurog::miniSOG2-CAAX and pSP-Neurog::mCherry-CAAX were ablated from the lateral side by irradiation with a laser (440 nm: 15 μW/cm^2^) for 10 min (Fig. 4A).

For the control, embryos electroporated with pSP-Neurog::mCherry-CAAX were ablated from the lateral side by irradiation with a laser (440 nm: 15 μW/cm ^2^) for 10 min (Fig. 4A). The ROI (region of interest) for laser irradiation was set to the region of mCherry fluorescence. Ablation was performed once per minute to correct the position of the ROI. After laser irradiation, the behaviour of embryos was recorded using a high-speed camera (WRAYCAM-VEX230M, WRAYMER, Osaka, Japan) attached to a stereoscopic microscope (OLYMPUS, SZX12, Tokyo, Japan) every 2 min for 30 min. Optical images were captured at 200–500 fps to quantify the temporal change in curvature associated with early spontaneous motor behaviour.

### Quantification of the ETF

To quantify early spontaneous motor behaviour, recording data were processed using the open-source program Fiji (National Institutes of Health, Bethesda, MD, USA) and MATLAB (MathWorks). The binarisation and skeletonisation ^40^ of the images were processed by Fiji, and then skeleton pruning was processed using a custom-made MATLAB script (MathWorks). The midline of the embryos at each time point was extracted, and the curvature of the midline was calculated using the position of three points on the midline: start, (A), mid-(B), and end (C). The equation for calculation of curvature was as follows.

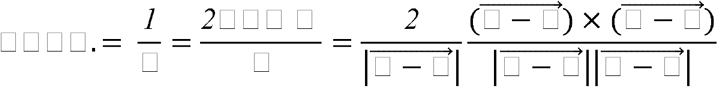

## Acknowledgments

*Ciona intestinalis* type A was provided by Dr. Yutaka Satou at Kyoto University and Dr. Manabu Yoshida at The University of Tokyo, with support from the National Bio-Resource Project of AMED, Japan. Valuable discussion and comments were provided by Dr. Atsuo Nishino.

This work was supported by JSPS KAKENHI Grants 16H01451, 16K07426 and 21H00440 (to K.H.), 19J21665 (to T.A.), 19H03213 (to T.G.K.), 17K15130 (to K.Oo.), and 19H03204 (to T.H.). K.H. is also supported by Keio Gijuku Academic Development Funds. T.H. is also supported by a Toray Science and Technology Grant, the Takeda Science Foundation, and the Mochida Memorial Foundation for Medical and Pharmaceutical Research. K.Oo. is also supported by a Sasakawa Scientific Research Grant from The Japan Science Society.

## Author Contributions

K.H. conceived this study; T.A., K.Ok., and K.H. designed the research; T.A. performed the research; K.Oo., M.M., T.H., T.G.K., and K.H. contributed reagents and analytic tools; K.Oo., M.M., and T.G.K. analysed cell lineage; T.A. analysed the data; T.A. and K.H. wrote this manuscript and designed figures; and K.Ok. and K.H. critically revised the manuscript and supervised the research.

